# Chronic wounds and adaptive *Pseudomonas aeruginosa*: A phenotypic and genotypic characterization

**DOI:** 10.1101/2024.07.08.602036

**Authors:** Kasandra Buchholtz, Rie Jønsson, Rasmus L. Marvig, Biljana Mojsoska, Karen Angeliki Krogfelt

## Abstract

Phenotypic and genetic diversity is found in varying prevalence in clinical populations where beneficial adaptations enable the bacteria to avoid recognition and eradication by the host immune system. This study aimed to investigate the presence of *Pseudomonas aeruginosa* in chronic venous leg ulcers wounds over an 8-week time course. This was performed using genomic and phenotypic approaches to understand the survival and persistence of *Pseudomonas* strains. The findings of this study show that the two patients were colonized with a recurring *P. aeruginosa* genotype with only minor phenotypic differences and few SNP differences, suggesting that the *Pseudomonas* isolates present in the wound can survive and proliferate in the host’s hostile environment. The results provided from this study will allow us to understand clinically important traits in *P. aeruginosa* and provide insight into the dynamics and adaptations (or the lack thereof) in chronic wounds.

**Highlights:**

- Each patient is colonised overtime with a unique MLST
- The total number of detected SNPs was 18 and 20 within each patient, respectively
- One MLST belonged to the sequence type ST132, often observed in clinical isolates, whereas the other belonged to ST3244, previously seen inenvironmental isolates

## 1. Introduction

The Gram-negative bacterium *Pseudomonas aeruginosa* is an opportunistic human pathogen. Responsible for infecting immunocompromised patients, *P. aeruginosa* is often seen in individuals with cystic fibrosis, urinary infections, and chronic wounds (Moradali et al., 2017). Wounds infected with *P. aeruginosa* are important in diabetic patients as they heal very slowly, and in venous leg ulcers are seen to be larger in size and associated with more severe wound outcomes compared to wounds where the pathogen is not present (Gjødsbøl et al., 2006; Halbert et al., 1992; Kirketerp-Møller et al., 2008; Madsen et al., 1996).

The large genome size and the genetic complexity of *P. aeruginosa* could explain the bacterium’s pathogenicity and unique ability to persist and thrive under different environmental conditions (Stover et al., 2000). In chronic wounds, the expression of multiple virulence factors (such as pyocyanin) is regulated by quorum sensing systems such as *las, rhl,* and *pqs* (Muller et al., 2009; Serra et al., 2015). Pyocyanin is a redox-active phenazine pigment that can interact with molecular oxygen and induce oxidative stress and cell damage to the host. This release of pyocyanin has been shown to enhance the adhesion, microcolony, and biofilm formation of *P. aeruginosa*, thus promoting the ability of the bacterium to form chronic wound infections. (Hall et al., 2016).

Treating *P. aeruginosa* infection is extremely difficult once established, probably due to the formation of antimicrobial-tolerant biofilms (Olivares et al., 2020). Therefore, studies are needed to investigate *P. aeruginosa* in clinically relevant environments to find novel ways of eradicating chronic infections.

This study characterized 16 clinical wound isolates of *P. aeruginosa* collected from two patients with chronic venous leg ulcers over an 8-week course for their phenotypic and genotypic diversity. We aimed to identify the clonal lineages to explore the hypothesis of whether the clinical wound isolates originated from the same hospital lineage of *P. aeruginosa.* Furthermore, we compared the strains phenotypically and genotypically to characterize the heterogeneity among the isolates during a chronic infection.

## 2. Materials and methods

### 2.1. Bacterial strains and growth conditions

Clinical wound isolates were collected over two months from patients admitted with persistent venous leg ulcers at Copenhagen Wound Healing Centre, Bispebjerg Hospital in Denmark (Gjødsbøl et al., 2006). Isolates were collected using filter paper pads, biopsies, and charcoal swabs throughout eight weeks of regular wound observations. Two patients (one male and one female) with repeated *P. aeruginosa* infection were chosen (Table 1). None of the patients had received antibiotics at least 14 days before or during the study period (Gjødsbøl et al., 2006). All samples were cultured, and bacterial isolates were stored at –80 °C in Luria-Broth (LB) with 10 % glycerol. All strains were grown at 37 °C. The clinical isolates used are listed in Table 2. Reference strains *P. aeruginosa* PAO1 and PA14 were used for comparison (He et al., 2004; Stover et al., 2000).

**Table 1.** Patient characteristics.

|  | Sex | Age | Duration of infection prior to inclusion (months) | Wound size (cm<sup>2</sup>) week 1 | Wound size (cm<sup>2</sup>) week 8 | % difference in wound size In the 8 weeks | Organisms identified besides *P. aeruginosa* |
| --- | --- | --- | --- | --- | --- | --- | --- |
| Patient id.3 | Male | 87 | 36 | 19.67 | 29.62 | 2.39 | *E. faecalis* |
| Patient id.5 | Female | 81 | 50 | 98.34 | 79.55 | -6.37 | *Yeast, S. maltophilia, E. cloacae, E. coli, E. faecalis, Moraxella spp.* |

**Table 2.**
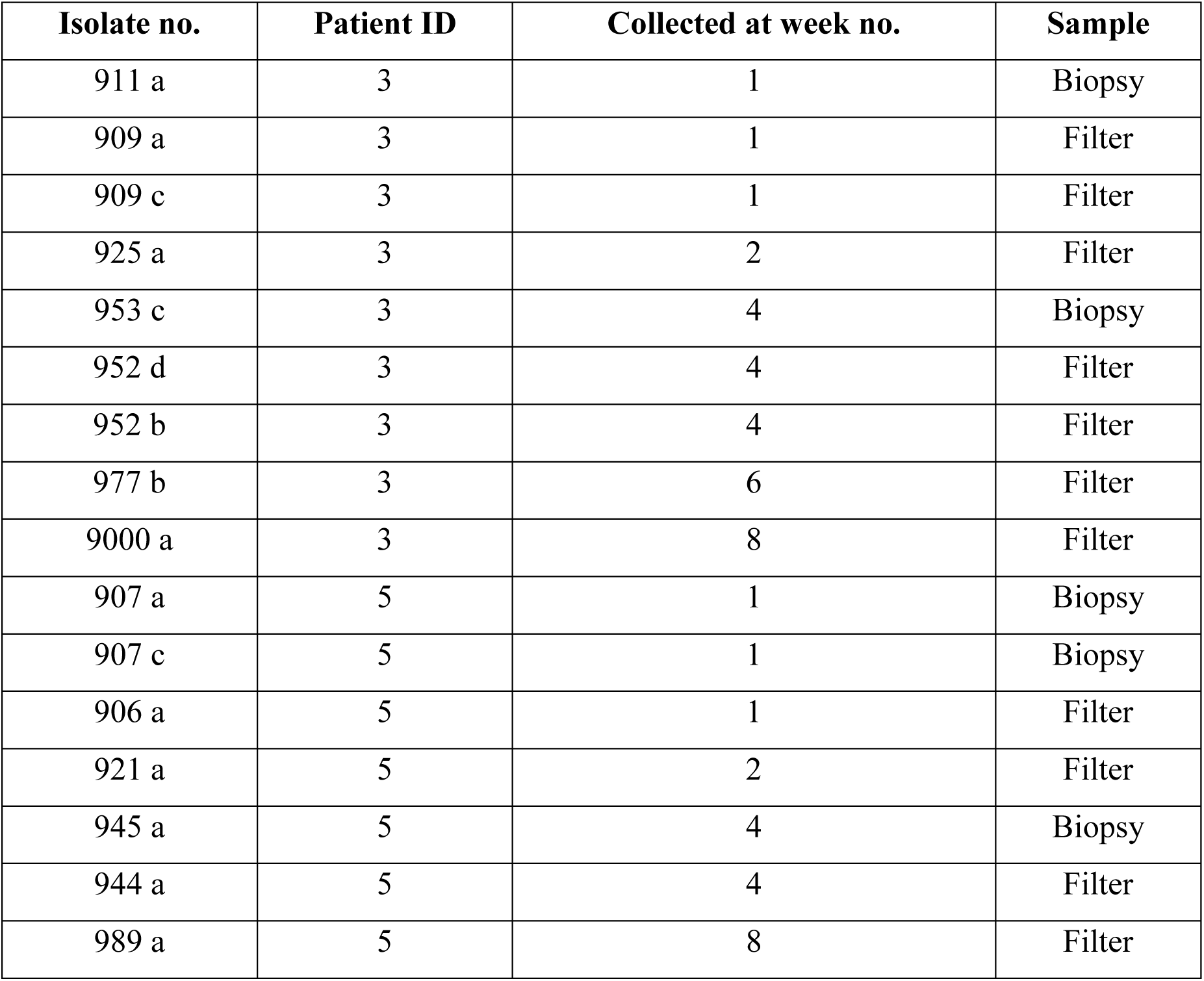
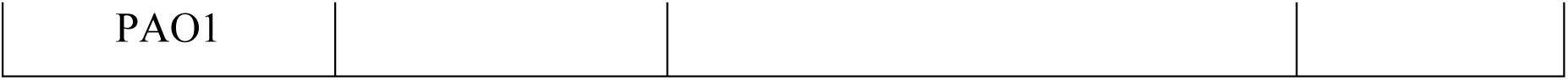
Clinical wound isolates were used in this study.

### 2.2. Growth rate determination

Overnight cultures diluted in LB-broth (Merck, Denmark) to OD_600_ 0.03 were used for growth rate measurements in a conical flask. All strains’ growth rates were measured by measuring the optical density during growth (∼20min) in 50 ml LB-broth in 250-ml flasks at 180 RPM. Growth rates were expressed as generation times in minutes. Three biological replicas for each strain were made.

### 2.3. Quantification of pyocyanin

A serial dilution of pyocyanin in LB-broth (100, 50, 25, 12, 6 µM) was made from 1 mg/ml of pyocyanin as a reference. UV-vis spectrophotometry was used to quantify pyocyanin in LB media. One colony of each clinical isolate of *P. aeruginosa* and PAO1 was inoculated in 5 ml of LB-broth and incubated overnight (O.N.) in a water bath at 37 °C, 180 RPM. After that, bacterial supernatants were filtered using a 0.45 µm filter (Frisenette Aps). The production of the quorum-sensing pigment was determined by measuring the absorption at 311 nm. The absorption at 311 nm was converted into a concentration using pyocyanin’s extinction coefficient in LB media. For normalization, a colony forming units (CFU) assay was performed from the overnight culture.

### 2.4. Determination of the minimum inhibition concentration

Clinical relevant antimicrobials were used to determine the phenotypic antimicrobial susceptibility (tobramycin, ciprofloxacin, meropenem, and polymyxin B) (n=3). A cation-adjusted Mueller-Hinton Borth (MHIIB) (Merck, Denmark) was used to test the susceptibility to tobramycin (Wiegand et al., 2008). The concentration range of the antibiotics used in this study was as follows: tobramycin 32-0.05 µg/ml; ciprofloxacin 20-0.031 µg/ml; meropenem, and polymyxin B, 40-0.062 µg/ml. The plate was sealed and incubated O.N. at 37 °C. CFU assay was performed on the 1:500 diluted culture to estimate the final concentration. The expected concentration range was 2×10^5^ – 8×10^5^ CFU/ml. The results were obtained 24 hours after incubation.

### 2.5. Swarming activity

The swarming motility of the *P. aeruginosa* strains was assessed as previously described (Song et al., 2019). Briefly, 5 µl O.N. cultures were plated from each culture in the center of a swarming plate, which contained M9 medium supplemented with MgSO_4_ (1mM), 0.5 % casamino acids, 0.2 % glucose and solidified with 0.5 % Bacto agar. Plates were incubated at 37 °C for 24 hours and plates were evaluated visually for swarming capabilities “+” or no swarming capabilities “-“. Each swarming experiment was carried out with at least three biological replicates.

### 2.6. Extraction of chromosomal DNA for genome sequencing

The extraction of genomic DNA was prepared according to a previously described protocol (Rasool et al., 2021) The concentration was measured on NanoDrop, and the samples were run on 0.7% agarose gel (Sigma) together with a Generuler 1 kb ladder (Thermo Scientific) to check for impurities. The sample was sent to Novogene for whole genome sequencing using the Illumina NovaSeq 6000 platform.

### 2.7. Genomic assembling

De novo assembly and annotation were created using the Shovill pipeline (https://github.com/tseemann/shovill). Next, BBDuk was used in the quality-trimming process. *In silico* analysis was conducted using ResFinder to identify resistance genes within the genome, and Multi-Locus sequence typing was performed using MLST 2.0. (Database version 24.04.2021)(Centre for Genomic Epidemiology (CGE)). Moreover, SNPs were identified in each isolate and compared across the isolate within the same sequence type using BacDist (https://github.com/MigleSur/BacDist). The BacDisk pipeline was set to filter and retain SNPs with coverage of at least ten reads in all clones and exclude mutation in clones with more than 80% non-reference reads. The reference genome used in this study was *Pseudomonas aeruginosa,* Stain: PAO1 (RefSeq assembly accession number: GCF_00006765.1). The Phylogenetic tree was visualized using the bioinformatic tool Tree of Life Tool – iTOL (version 6).

### 2.8. Single-nucleotide-variant (SNV)-based phylogenetic tree

We used parnsp version 1.2 (Treangen et al., 2014) with default settings to make core genome single-nucleotide-variant (SNV)-based phylogenetic tree of assembled genomes.

### 2.9. Statistics

Statistical analyses were performed in GraphPad Prism 9.3 (GraphPad Software Inc., CA, USA). Doubling times were compared by a one-way ANOVA and Tukey’s multiple-comparisons test. P value below 0.05 (p < 0.05) was considered statistically significant. Further levels of significance are indicated with asterisks *P < 0.05, ** P < 0.01, *** P < 0.001.

### 2.10. Ethics statement

The Danish Scientific Ethical Board approved the study, and samples were obtained after all patients’ written consent.

## 3. Results

To investigate *P. aeruginosa* diversity during chronic wound colonization, isolates from two patients (patient id. 3 and patient id. 5) with repeated *P. aeruginosa* infection were chosen. Table 1 summarizes the patient characteristics.

### 3.1. No differences in growth rates between the isolated strains

Long-term infection of *P. aeruginosa* can sometimes be accompanied by a reduction in growth rate for a pathogen to adapt to its environment (la Rosa et al., 2021). This was tested by measuring the growth rate of clinical wound isolates and comparing it to the reference strain PAO1. The linear regression result showed the doubling time for the clinical isolate from patient id. 3 had a doubling time that varied from 39 to 45 minutes, whereas for patient id. 5, the doubling time varied from 37 to 55 minutes (Table 3). In comparison, PAO1 had a doubling time of 37 minutes. A one-way ANOVA was used to compare each patient’s wound isolates. The result showed no significant difference in the doubling time between the isolates from patient id. 3 nor patient id. 5.

**Table 3.**
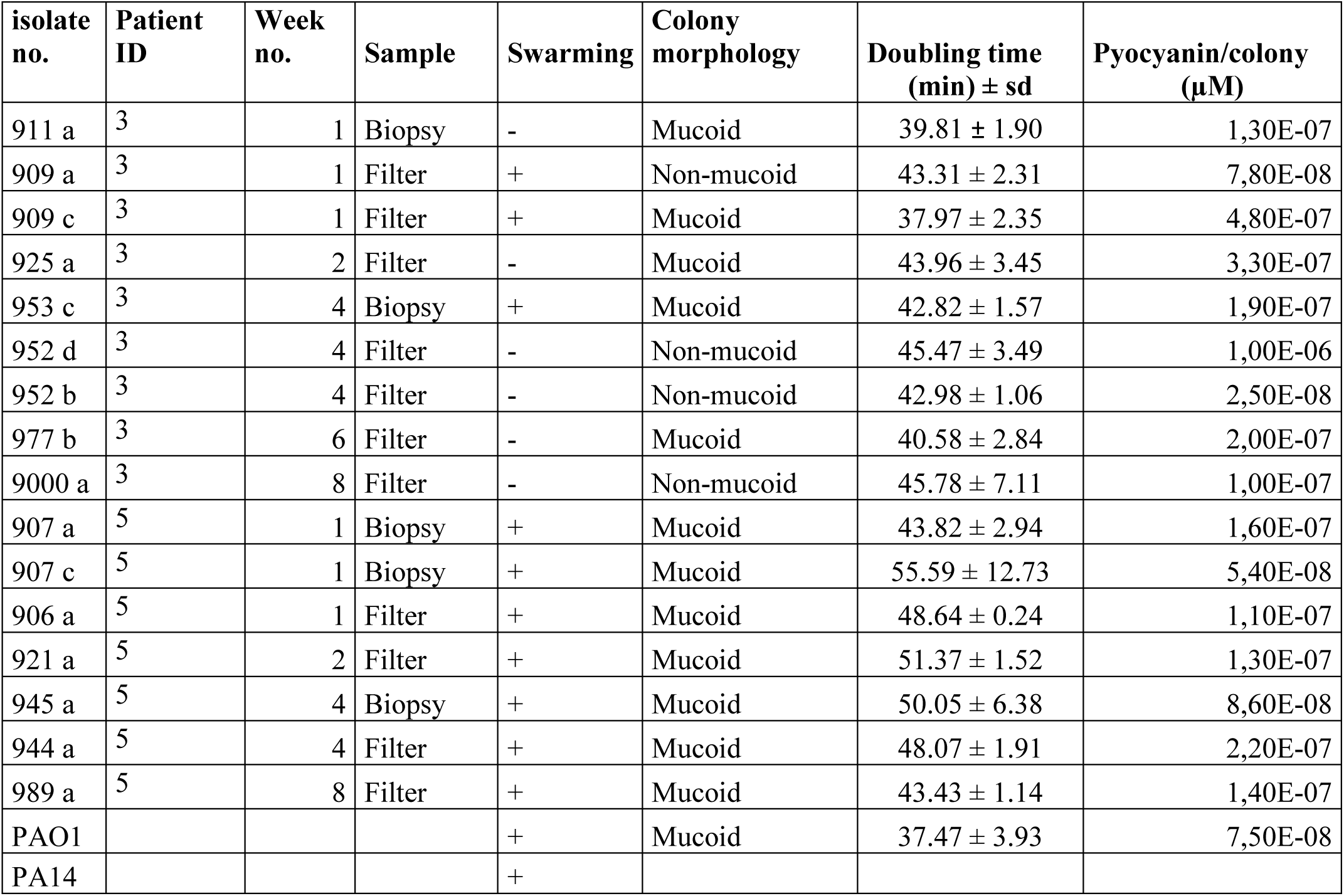
Summary of the phenotypic analysis performed on *Pseudomonas aeruginosa* wound isolates from patients id 3 and 5 and reference strain PAO1 and PA14. In the colony morphology analysis, they were distinguished between mucoid and non-mucoid. The doubling time is minutes (± SD; n = 3). Swarming was evaluated visually, where ‘+’ indicates swarming capabilities, and ‘-‘ is when no swarming is observed. In the quantification of quorum sensing pigment, pyocyanin was used as a reference molecule, and the amount of pigment produced is given in µM/CFU ml^-1^

### 3.2. Small phenotypic differences were observed in patient 3 for both swarming and colony morphology

*P. aeruginosa* harbors polar flagella, which enables it to swarm on semi-solid agar. This complex behavior allows it to move across a surface and is suggested to play a role in the initiation of biofilm formation (Overhage et al., 2008; Shrout et al., 2006; Yeung et al., 2009). To assess the swarming abilities of the strains, overnight cultures were spotted on 0.5% M9 agar plates supplemented with casamino acids and glucose. The results showed that six of the strains from patient id. 3 were not able to swarm, except for three strains (909a (week 1), 909c (week 1) and 953c (week 4) (Figure 1). In patient id. 5, all strains could swarm. There was no correlation between the colony phenotype and the isolation method, i.e., filter and biopsy. The colony morphology analysis observed that the wound isolates 952d and 909a (weeks 1 and 3) from patient id. 3 were non-mucoid, whereas all the other clinical wound isolates from patient id. 3 and id. 5 and were mucoid, together with PAO1 (Table 3).

**Figure 1.**
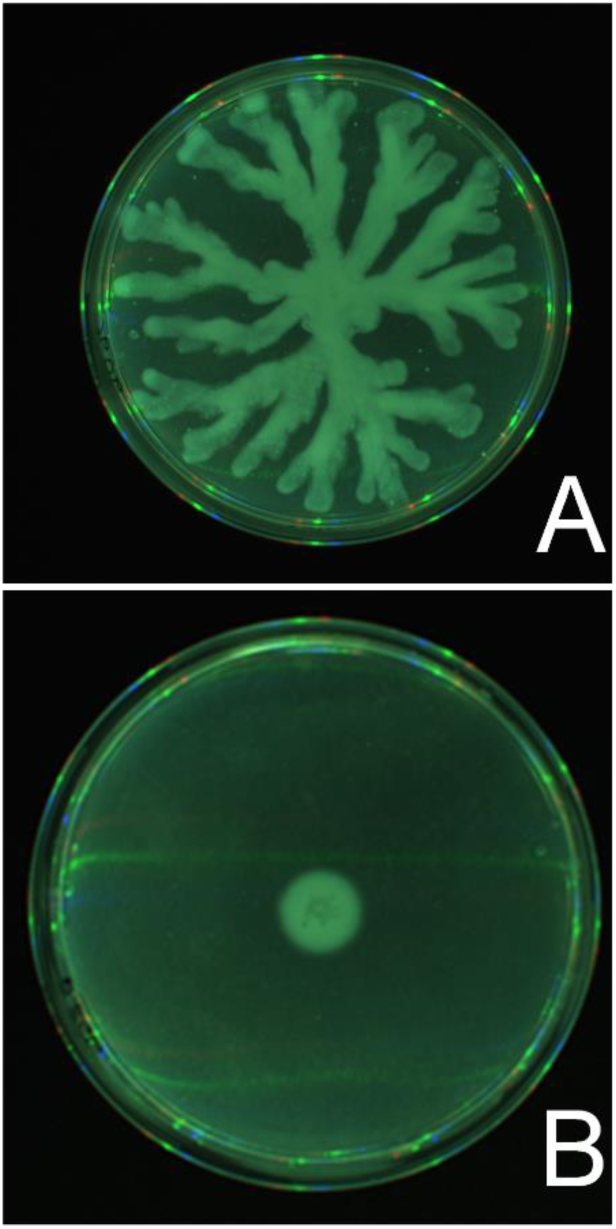
Some isolates were capable of swarming in patient id. 3 (isolate 909c, week 1) whereas other isolates from the same patient (e.g. 952d, week 4) showed no swarming capabilities.

### 3.4. Strain 953c (week 4) produces significantly more pyocyanin compared to the other strains

To investigate the phenotypic diversity of the clinical wound isolates *of P. aeruginosa* and PAO1, spectrophotometric analysis was performed to estimate the amount of quorum-sensing pigment pyocyanin produced by each clinical isolate and PAO1. The result of ANOVA analysis followed post hoc Turkeys multiple comparison tests showed a significant difference between isolate 953c and isolate 911a, 925d, 952b and 977b (** p < 0.01) for patient id. 3 (Figure 2). The test results for patient id. 5 showed no significant difference between the isolates.

**Figure 2.**
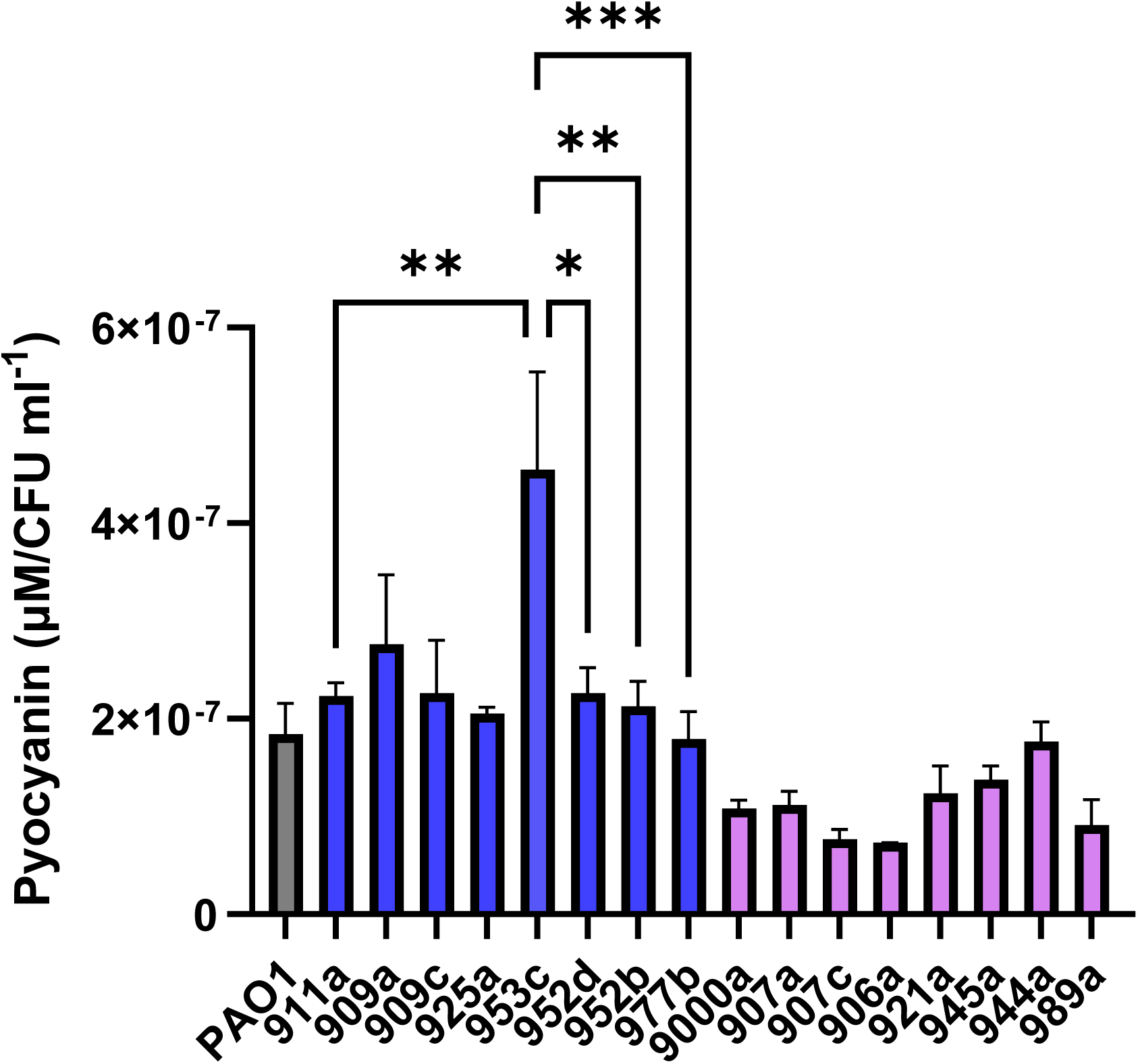
Graphic representation of the quorum sensing pigment produced by each clinical isolate of *Pseudomonas aeruginosa* from patient id. 3 (blue) and id. 5 (purple) together with PAO1 (Gray)., The data was obtained using UV-Vis spectroscopy using Pyocyanin as a reference molecule and 311 nm absorption wavelength.Thethe result was normalized to represent the concentration µM per CFU/ml. (± SEM; n=3).

### 3.5. The patients carried the same MLST type during the 8 weeks, yet different MLSTs

The phylogenetic diversity among the isolates was determined by measuring the single nucleotide polymorphism (SNPs) present in each genome, thereby revealing the evolutionary relationship between the wound isolates. The MLST analysis showed that the seven wound isolates were collected from patient id. 3 had the same sequence type profile ST132. The same observations were seen for the nine wound isolates collected from patient id. 5, with the sequence type profile ST3244, and for PAO1, the sequence type profile was ST549 (Figure 3).

**Figure 3.**
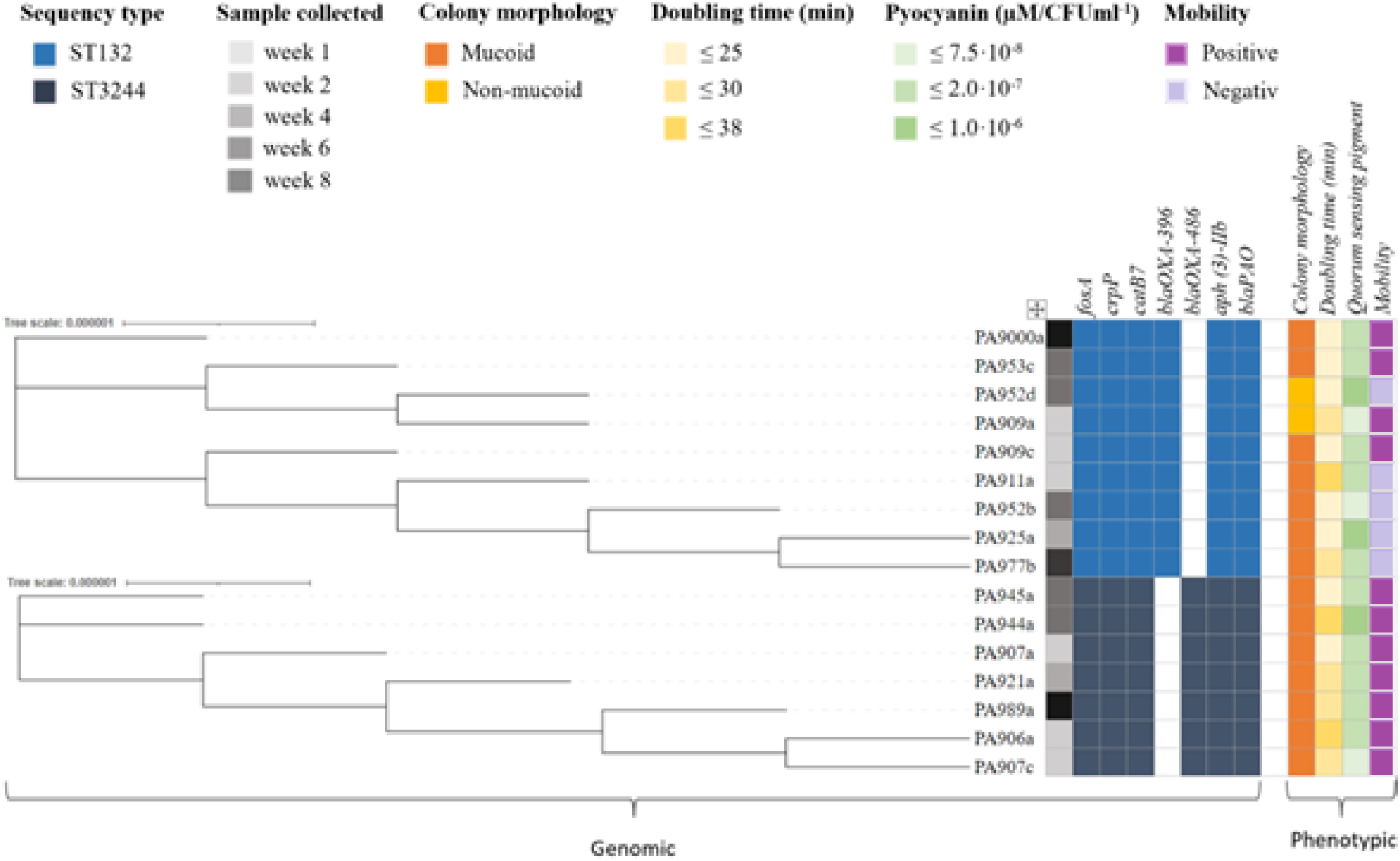
Phylogenetic tree, antibiotic resistance genes, and the phenotypic profiles of the strains. The phylogenetic tree was generated by analyzing the single nucleotide polymorphisms (SNPs). The tree scale indicates the substitution per site.

The total number of SNPs found in ST132 (n = 9) was 18; in ST3244 (n = 7), the total number of SNPs was 20. Moreover, the phylogeny results showed a close genetic relationship between all ST132 wound isolates, with 0-8 SNPs differences (median 4 SNPs differences). The same minor SNPs difference was observed between all ST3244 wound isolates, with 0-15 SNPs differences (median 7 SNPs differences). The phylogenetic diversity is illustrated in Figure 3, together with the antibiotic resistance profile found in the ST132 isolate and ST3244 isolate. When we aligned the genomes of our 17 isolates to a collection of 448 *P. aeruginosa* genomes (446 genomes from Gabrielaite et al. (Gabrielaite et al., 2020) (and genomes of reference strains PAO1 (NC_002516.2) and PA14 (NC_008463.1)), we found that genomes of ST132 were within PAO1 main group, whereas the genomes of ST3244 were not part of either PAO1 or PA14 main groups (Figure 4).

**Figure 4.**
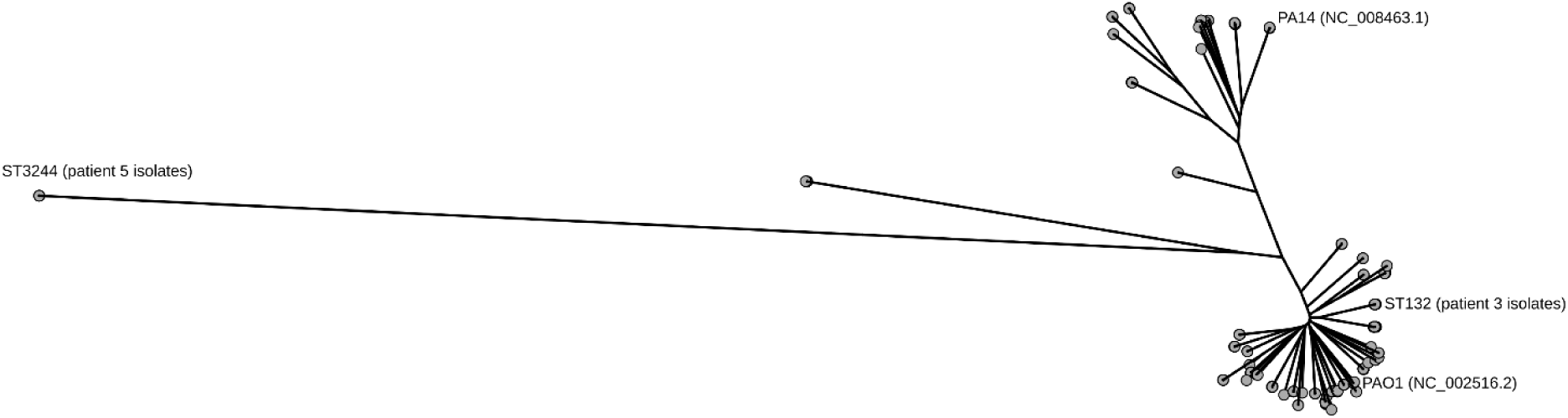
Phylogenetic tree of *P. aeruginosa* isolates. The phylogenetic tree was reconstructed using 13,728 polymorphic nucleotide sites and rooted with two *B. abortus* strains.

### 3.6. Intralineage genetic diversity was observed in the two proteins PilA and AmrZ

To investigate whether intra-lineage genetic diversity was occurring in the chronic wound, DNA sequence analysis was performed on 12 essential proteins for colonization, virulence, and biofilm formation (PilA, PilU, PilT, PilR, FliC, AmrZ, RhlR, LasR, FlgK, MucA, AlgU, AlgR). Among the mentioned genes, we only observed intralineage mutations in two proteins (PilA and AmrZ) within some of the isolates from patients’ id. 3 (table 4).

**Table 4.**
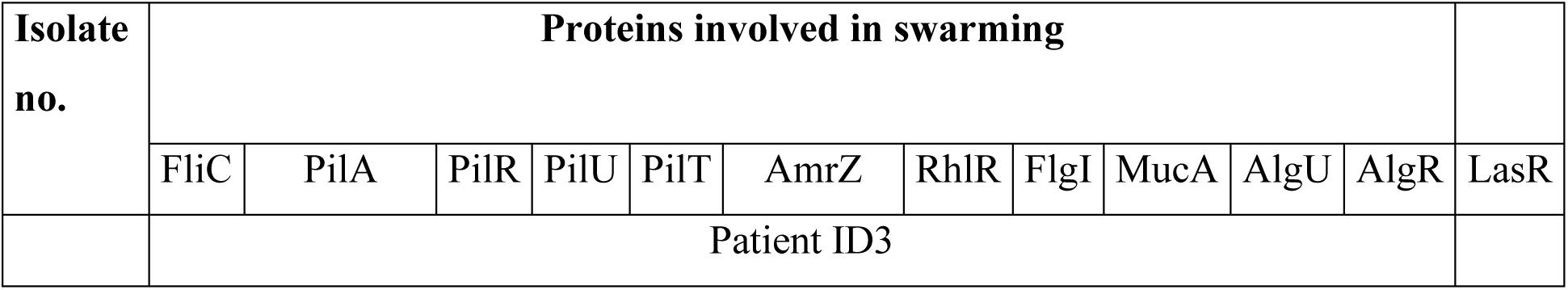
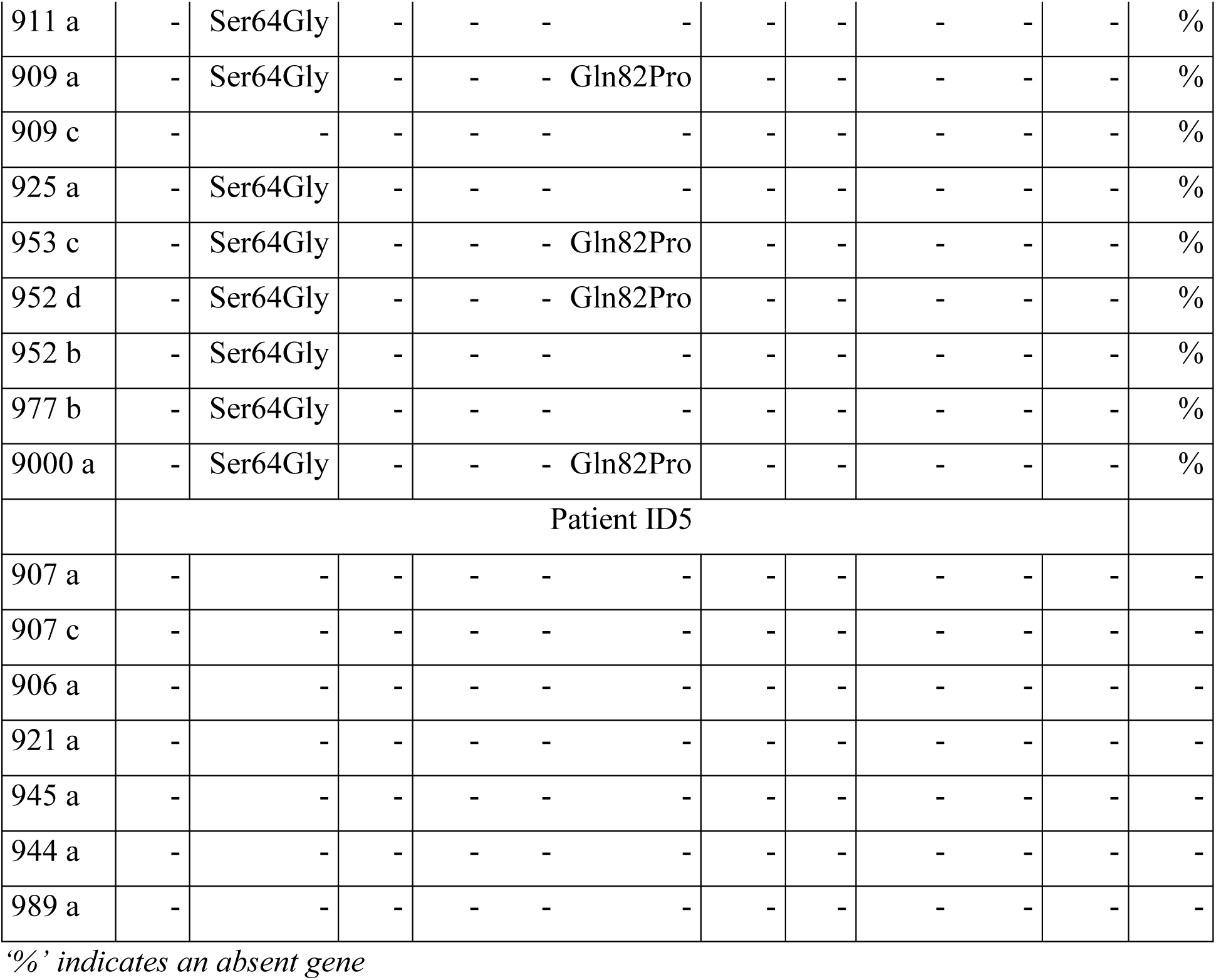
Intralineage mutations in swarming proteins.

**Table 5.** The minimum inhibition concentration was determined with the use of tobramycin (3.2-0.005 µg/ml), Meropenem (4-0.0062 µg/ml), ciprofloxacin (2-0.0031 µg/ml), and Polymyxin B (4-0.0062 µg/ml). The MIC breakpoint for *P. aeruginosa*, according to European Committee on Antimicrobial Susceptibility Testing (EUCAST), is for tobramycin (S ≤ 2; R > 2), meropenem (S ≤ 2; R > 8), ciprofloxacin (S ≤ 0.001; R > 0.5) and for Polymyxin B (S ≤ 2; R > 2). The capital letter S and R stand for susceptibility and resistance, respectively.

| Isolate no. | Tobramycin | Meropenem | Ciprofloxacin | Polymyxin B |
| --- | --- | --- | --- | --- |
|  | MIC (µg/mL) |  |  |  |
| Patient ID3 |  |  |  |  |
| 911 a | 0,4 | 20 | 0,25 | 1\|0.5 |
| 909 a | 0,4 | 0,025 | 0,25 | 0,1 |
| 909 c | 0,4 | 0,05 | 0,25 | 0,1 |
| 925 a | 0,4 | 0,5 | 0,25 | 0,5 |
| 953 c | 0,4 | 0,05 | 0,25 | 0,2 |
| 952 d | 0,4 | 0,05 | 0,25 | 0,2 |
| 952 b | 0,4 | 0,05 | 0,25 | 0.2\|0.1 |
| 977 b | 0,4 | 0,05 | 0,25 | 0,1 |
| 9000 a | 0,4 | 2 | 0,25 | 4\|2\|1 |
| Patient ID5 |  |  |  |  |
| 907 a | 0,8 | 0,05 | 0,5 | 0,1 |
| 907 c | 0,4 | 0,025 | 0,25 | 0,1 |
| 906 a | 0,4 | 2 | 0,25 | 2 |
| 921 a | 0,4 | 0,05 | 0,5 | 0,1 |
| 945 a | 0,4 | 2 | 0,5 | 0,5 |
| 944 a | 0,4 | 0,1 | 0.25\|1 | 0,1 |
| 989 a | 0,4 | 2 | 0,25 | 1 |
| Reference strain |  |  |  |  |
| PAO1 | 0,4 | 20 | 2 | 0,25 |

### 3.7. No phenotypic antimicrobial resistance against the antibiotics tested

The results from the bioinformatic analysis revealed that all wound isolates harbored resistance genes. This included fosfomycin (*fosA)*, ciprofloxacin (*crpP)*, chloramphenicol (*cat7B*), a carbapenem (*bla_OXA-396_*/*bla_OXA-486_*), aminoglycoside (*aph(3’)-IIb*), and β-lactam (*bla_POA_*). However, when performing minimum inhibitory concentration testing on the strains, the strains showed all to be susceptible to common antibiotics used for *P. aeruginosa* infections, such as tobramycin (aminoglycoside), ciprofloxacin (fluoroquinolone), meropenem (carbapenem (β-lactam)), and polymyxin B (table 4).

## Discussion

Residing in a laboratory culture flask is a stark contrast to inhabiting a chronic wound, where bacteria endure ongoing environmental stressors, including challenges from the host’s immune system, pH levels, temperature fluctuations, bacterial diversity, and limited access to nutrients and oxygen. These conditions are believed to prompt the development of adaptive strategies in bacteria to thrive in chronic infection. This study investigated the complexity and stability of *P. aeruginosa* populations in 2 chronic wounds during an 8-week course. Sequencing of the isolates revealed that each patient was colonized with different genotypes, and the isolates from each patient remained relatively stable during the 8 weeks.

Bacteria isolated from chronic infections are often thought to either lose the function of genes involved in virulence or at least alter their virulence (Darch et al., 2015; Mowat et al., 2011; Smith et al., 2006a; Winstanley et al., 2016). Patient id. 3 harbored the ST-type ST132, frequently observed in clinical settings, whereas ST3244 from patient id. 5 has only been reported in the environment. Notably, the two patients were treated in the same hospital setting and during the same period of type. This fact suggests that the patients were colonized by an isolate that was adapted to the patient itself. The total number of SNPs found in ST132 was 18, and in ST3244, the number was 20. In comparison, a study performed on *P. aeruginosa* in cystic fibrosis patients showed 41 SNPs over a 7.5 year period (Smith et al., 2006b), whereas another study showed 15 SNPs during 15 years (Cramer et al., 2011).

Previous studies have shown that diversity in a bacterial population *in vivo* is often related to biofilm formation, growth rate, and motility. Among growth rates, no difference was observed between the strains of patient id. 3 and id. 5. Interestingly, some of the isolates from each patient could not swarm on semi-solid agar, suggesting diversity among them. The ability to swarm is probably essential in the initial colonization, whereas when the pathogen has established itself in the chronic wound, swarming might not be as important. Differences within an isogenic population could result from variations in the cells’ ability to adapt to environmental changes (Vermeersch et al., 2019). This means that even though a difference in the swarming is observed between the wound isolates, they still originate from the same clonal lineage.

We observed some mutations in swarming-related genes (table 4), but other isolates who were swarming also harbored the same mutation. Previous studies (Overhage et al., 2008; Tremblay and Déziel, 2010) have shown that over 378 genes are involved in swarming motility, which suggests that deficiency in swarming motility can be challenging to identify genotypically. The virulence factor pyocyanin was produced by all isolates, though in higher amounts in the ST132 representing the clinical isolates. Yet, it has previously been suggested that a lysophospholipid (MPPA), which is generated by the secretory phospholipase A2 and accumulates in inflammatory exudates, inhibits the production of virulence factors by inhibiting the Quorum sensing system (Laux et al., 2002).

The study shows that the two patients were colonized with the same *P. aeruginosa* isolates over time with minor phenotype differences. Compared to the studies shown in cystic fibrosis lungs, adaptations in a chronic wound may be less extensive, as demonstrated by a study on a murine wound model (Vanderwoude et al., 2020). However, given the knowledge that the infection was between 36 and 50 months at the inclusion of this study, caution must be exercised since the *P. aeruginosa* strains can have colonized much prior to inclusion in this study.

In conclusion, our findings suggest that *P. aeruginosa* can colonize and persist in chronic wounds for extended periods. During these 8 weeks, only minor differences were observed between the isolates in the respective wounds. More studies are needed to better understand the diversity and evolutionary adaptations of *P. aeruginosa* during chronic wound infections.

## Author contributions

KB, RJ, RLM, BM, KAK: Conceptualization, Methodology, Software, Validation, Formal analysis.

KJP, RJ, RLM: Data curation, Writing original draft, Visualization, Investigation. RJ, KAK, BM: Reviewing and Editing.

All authors read and approved the final manuscript.

## Acknowledgment

Thanks to Kristine Gjødsbøl for providing culture data from the patient’s samples. Also, thanks to project nurses Lone Haase and Hanne Vogensen, Copenhagen Wound Healing Center, and Bispebjerg University Hospital for collecting samples and recruiting patients.

